# Estimates of local biodiversity change over time stand up to scrutiny

**DOI:** 10.1101/062133

**Authors:** Mark Vellend, Maria Dornelas, Lander Baeten, Robin Beauséjour, Carissa D. Brown, Pieter De Frenne, Sarah C. Elmendorf, Nicholas J. Gotelli, Faye Moyes, Isla H. Myers-Smith, Anne E. Magurran, Brian J. McGill, Hideyasu Shimadzu, Caya Sievers

## Abstract

Two recent meta-analyses of local-scale biodiversity change over time, by the authors of the present paper, have been subject to a harsh critique. Here we use new data and analyses to respond to the main points of this critique. First, a central argument of the critique was that short-term time series lead to biased estimates of long-term biodiversity change. However, we show here that this conclusion was based entirely on two fundamental mistakes in the simulations used to support it. Second, we show that the critic's conclusion that there are negative relationships between temporal biodiversity change and study duration is entirely dependent on: (i) the unrealistic assumption that biodiversity change can be positive when study duration = 0; (ii) the use of only a subset of the available data; (iii) inclusion of a single outlier data point in a single study (out of 100 in this case); and/or (iv) a choice to use log ratios rather than slopes (when available) as the metric of temporal biodiversity change. In short, the evidence does not support the conclusion that studies of longer duration tend to find local diversity decline. Finally, the critique highlighted the obviously true fact that studies in the ecological literature represent a geographically biased sample of locations on Earth; this issue was noted in both original papers, and is relevant for all ecological data syntheses. This fact was used by the critics to cast doubt on our conclusion that, outside of areas converted to croplands or asphalt, the distribution of temporal biodiversity trends is centered on zero. As a scientific rule, future studies based on more or different data may cause us to modify our conclusion, but at present, alternative conclusions based on the geographic-bias argument rely entirely on speculation. In sum, the critique raises points of uncertainty typical of all ecological studies, but it falls far short of providing an evidence-based alternative interpretation for our results, which are now supported by syntheses of hundreds of individual data sets of temporal biodiversity change.

## Introduction

Gonzalez et al. (in press) have raised concerns about two papers that collectively analyzed >250 individual datasets on biodiversity change through time from many parts of the world (Vellend et al. 2013, Dornelas et al. 2014). Both of these studies found that the average magnitude of temporal change across studies was indistinguishable from zero. The concerns of Gonzalez et al. are for the most part typical of those that could be directed at any ecological meta-analysis: different results might obtain in different places (underrepresented regions) or times (before people collected data of this nature), and it is possible to find data subsets that deviate from the overall pattern. These concerns were articulated by Gonzalez et al. in dramatic fashion so as to call into question our conclusions. Some aspects of the Gonzalez et al. critique were based on selective use of data and methods of analysis, while others focused on the nature of the data themselves and accompanying interpretations.

Here we present analyses, as well as new data, to support the following conclusions: (1) Shortterm time series do not provide biased estimates of long-term trends. The opposite conclusion presented by Gonzalez et al. was based on two basic errors in their simulation model and calculations. (2) There is no compelling evidence that studies of longer temporal duration show greater biodiversity decline. On this point, the analyses presented by Gonzalez et al. were fragile in the extreme with respect to single outlier data points, to assumptions about model structure, and to the inclusion of additional data. (3) The ecological literature is indeed geographically biased, a fact discussed explicitly in both Vellend et al. (2013) and Dornelas et al. (2014). The sophisticated analysis of Gonzalez et al. on this issue serves only to make the self-evident point that new data (in this case from underrepresented regions) might modify conclusions.

### (1) Short-term time series do not provide biased estimates of long-term trends

One key component of the Gonzalez et al. critique is in factual error (i.e., not a matter of selective interpretation). Simulations of species richness (S) over 50-year periods and subsequent calculations of log ratios (log(S_after_/S_before_)) or slopes of richness on time during shorter time intervals (5, 10, 20 years) were used to argue that “Estimates of biodiversity change are systematically biased when syntheses are based on datasets composed primarily of short time series”. Gonzalez et al. made two different errors, the first of which applies only to log ratios, the second of which applies to both log ratios and slopes:

i. When calculating a mean effect size for “short” windows of time, Gonzalez et al. did not take into account the fact that a log ratio for, let’s say, a 10-year period of time is only expected (mathematically) to capture one fifth of the amount of change that occurs over 50 years. In other words, they did not multiply the average of 10-year windows by 5 before comparing with the 50-year effect size. This is equivalent to the fallacious argument that, hypothetically, temperature only went up by 0.5C per decade, so the estimate of the “real” increase of 2.5C over 50 years is biased. This is obviously incorrect.
ii. The second problem is less obvious, but no less important, and it accounts for apparent diversity increases in medium-sized time windows (e.g., 20 years) when a 50-year period shows a richness decline initially, followed by an increase, and then a leveling off (see Fig. 1A-C). The problem is that with a bounded range of 50 years, “randomly” chosen segments of 20 years severely over-represent the middle portion of the time series. In another well-known ecological context, this is called the mid-domain effect to explain peak species richness at central latitudes or altitudes (Colwell and Lees 2000). However, whereas the boundaries in space are real, the temporal boundaries are not, as time is infinite in both directions. The first point in the time series, for example, is only part of one 20-year segment in the “population” from which the Gonzalez et al. simulations sample, 0:20. The second time point is part of two segments, 0:20 and 1:21, and so on. Time points 20-30, on the other hand, are each part of 20 different segments. So, with the decline in richness happening early during the 50-year time span, seemingly random samples of 20 years mostly miss the decline, while “detecting” a transient increase only because it happens to occur in the middle portion of the time series. The apparent bias detected by Gonzalez et al. is an artefact of their simulation analysis focusing on an arbitrary bounded time interval (Fig. 1).

**Figure 1.**
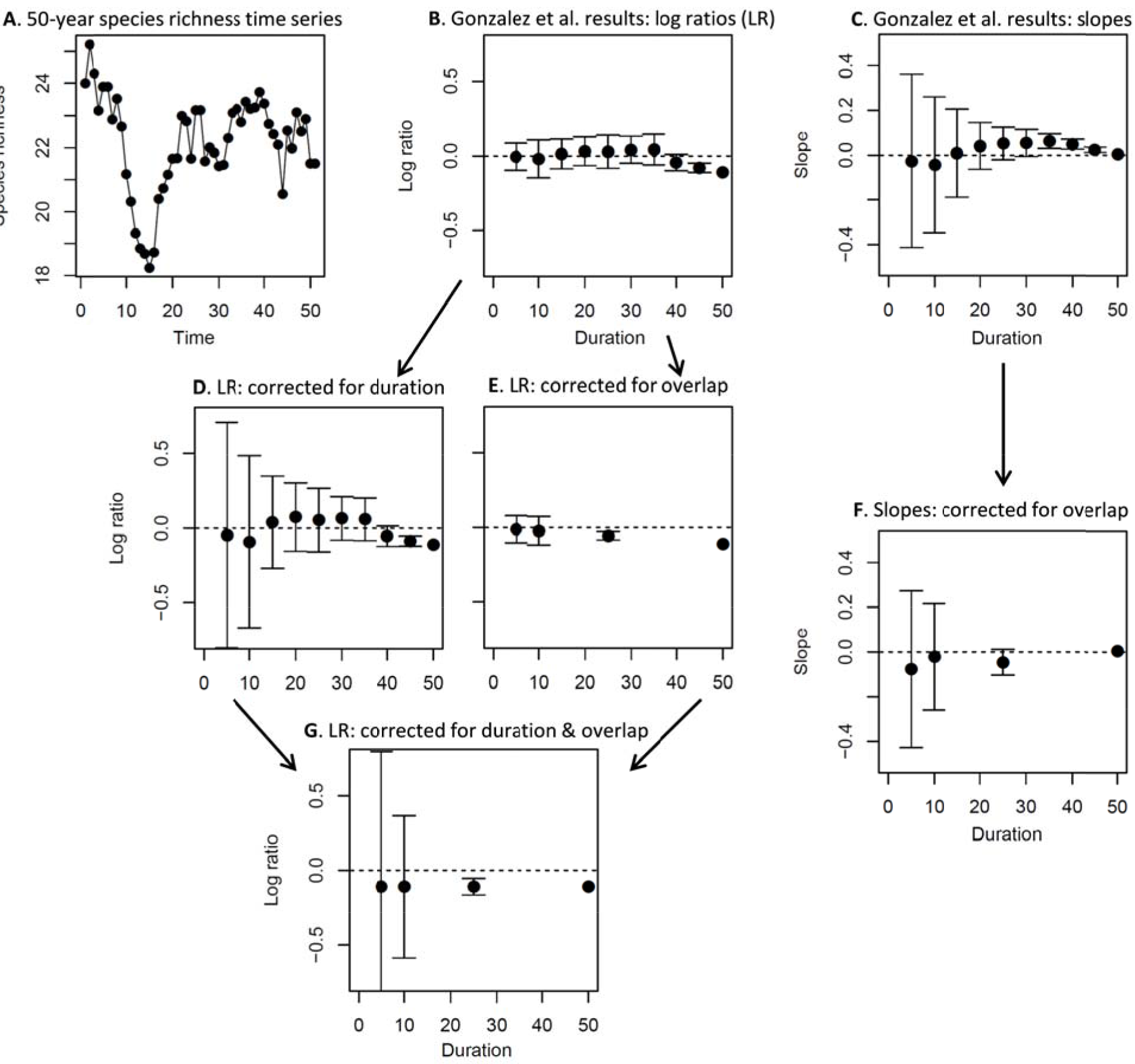
Mean ± standard deviation of log ratios (log(S_after_/S_before_)) and slopes (species richness vs. time) for repeated samples of short time series sampled from a longer (50-year) duration data set. **A**: A reproduction of Fig. S3D from Gonzalez et al., showing one example of species richness dynamics over time that appeared to lead to especially biased results. **B, C**: Results of 1000 seemingly random samples of different duration conducted according to the methods of Gonzalez et al.; these results appear to show an average positive trend among moderate-duration samples, despite a long-term negative (log ratio) or flat (slope) trend over the full duration. **D, E**: Log ratio results when correcting separately for duration (problem (i) in main text) and overlap (problem (ii) in main text); here we see that just accounting for the duration of data subsets removes bias from short-duration samples, while correcting for overlap removes any tendency for positive average trends. **F**: Slope results after correcting the overlap problem. **G**. Log ratio results after correcting for both problems; here the averages are precisely equal to the long-term trend. Note that when correcting for overlap, we only use durations that are multiples of the 50-year total time span.

If one examines sequential, non-overlapping portions of any length of a given time series, the average log ratio captures precisely the rate of change over the entire time series. Simulations are not required to demonstrate this point, although we provide one corrected example from Gonzalez et al. (Fig. 1), in addition to the following explanation from first principles. Imagine we have a species-richness (S) time series of five points, t_0_:t_4_, and thus four year-to-year transitions. The log ratio from beginning to end is log(S_4_/S_0_). The average of one-year intervals is:

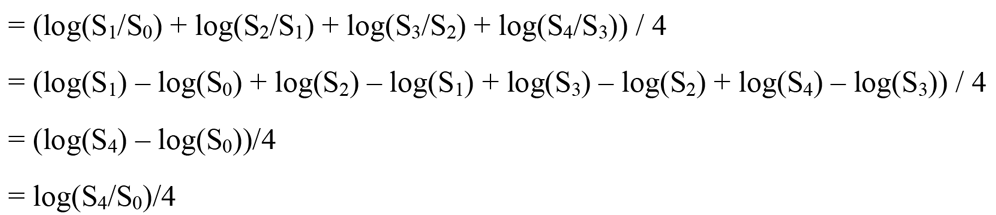

So, as long as we account for the fact that the one-year intervals cover only one quarter of the full time series (i.e., we multiply this by four), we recover the original “target” log ratio for the full time series precisely (see also Fig. 1G). The same result will hold for two-year intervals in this time series, 10-year intervals of a 50-year time series, or any other combination. The same precise mathematical equivalence does not hold for slopes, but it is equally true that there is no systematic bias introduced by the fact of sampling a subset of a longer time series. An incomplete sample of the portions of the longer time series will introduce variance (as is always the case with sampling), but not systematic bias (Fig. 1). The conclusion, based on simulations, “that short time series can provide unreliable estimates of a known trend” (Gonzalez et al. in press) is simply incorrect.

### (2) Local biodiversity trends in studies of different duration

Gonzalez et al. used their incorrect argument that short-term time series bias estimates of temporal biodiversity trends as a springboard to asking whether longer duration studies tend to show biodiversity declines. In this section, we address this issue for the two original studies in turn.

Using the data from Vellend et al. (2013), Gonzalez et al. modeled the log ratio of species richness at the end and start of a study (see previous section) as a function of the duration of that study, finding a statistically significant (p = 0.04) but weak relationship (Fig. 2A). They emphasized the conclusion that longer-duration studies tend to show richness declines, although by allowing for a non-zero intercept, their results also require explaining a nonsensical positive biodiversity trend in studies that last zero years. If one makes the ecologically realistic assumption that the log ratio must be zero at duration = 0 (i.e., a zero intercept), not only is the slope not significant, but its raw value is actually positive rather than negative (Fig. 2B). This illustrates the potentially major influence of assumptions about model structure on the spurious detection of weak statistical relationships.

**Figure 2.**
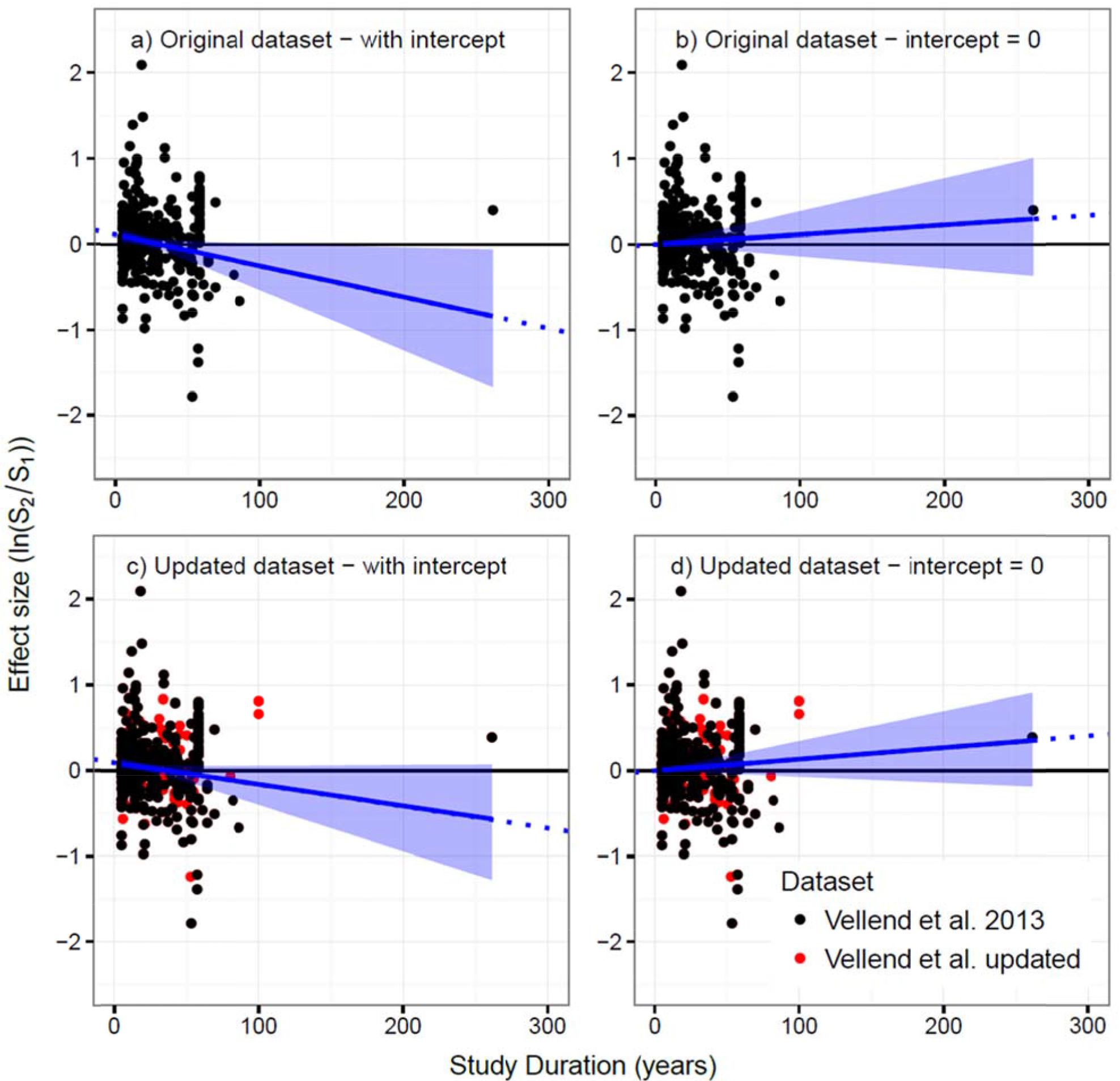
Relationships between local plant species richness change over time (y-axis) and the duration of a study, modeled assuming either a non-zero y-intercept (a,c) or a zero y-intercept (b,c), using the original data in Vellend et al. (2013) (a,b) or an expanded dataset (c,d; new data points shown in red). The effect size for temporal richness change is expressed as the log ratio of species richness in the final year of study (SR_2_) and in the initial year of study (SR_1_). Lines represent the estimated effect size with credible intervals. See Appendix A for statistical methods.

Given the controversy sparked by Vellend et al. (2013), we have since expanded the data set by 37% to include studies published through the end of 2014 (the original paper had studies published up to July 2012; https://github.com/lbaeten/DivChangeResponse). The methods were identical to those in Vellend et al. (2013), except that we did not additionally read through the references of all new papers to find additional data sets. With the larger data set of 212 studies (the 2013 paper had 155), there is no significant relationship between local richness change and study duration, regardless of whether one allows for a non-zero intercept (Fig. 2C,D).

The data in Dornelas et al. (2014) includes studies with diversity estimates for at least three time points, thus allowing the estimation of slopes of diversity vs. time, rather than only before-after log ratios. There is no significant relationship between the diversity-time slope and study duration (Fig. 3A,B). Gonzalez et al. chose instead to calculate log ratios using the data in Dornelas et al. (2014; see Dataset S1 in that paper), and reported a significant negative relationship between log ratios and study duration (Fig. 3C). Again their analysis allowed for a non-zero intercept, and if the intercept is fixed at zero the relationship is not significant (Fig. 3D). In addition, the Gonzalez et al. result is highly sensitive to one outlier, depending not only on a single study (reference 90 in Dornelas et al. 2014), but on a single data point in that study (species richness = 43 in 1911, and <20 for the next 90 years). In the absence of that one data point, the relationship is not statistically significant, regardless of whether one assumes a zero or non-zero intercept (Fig. 3E,F).

**Figure 3.**
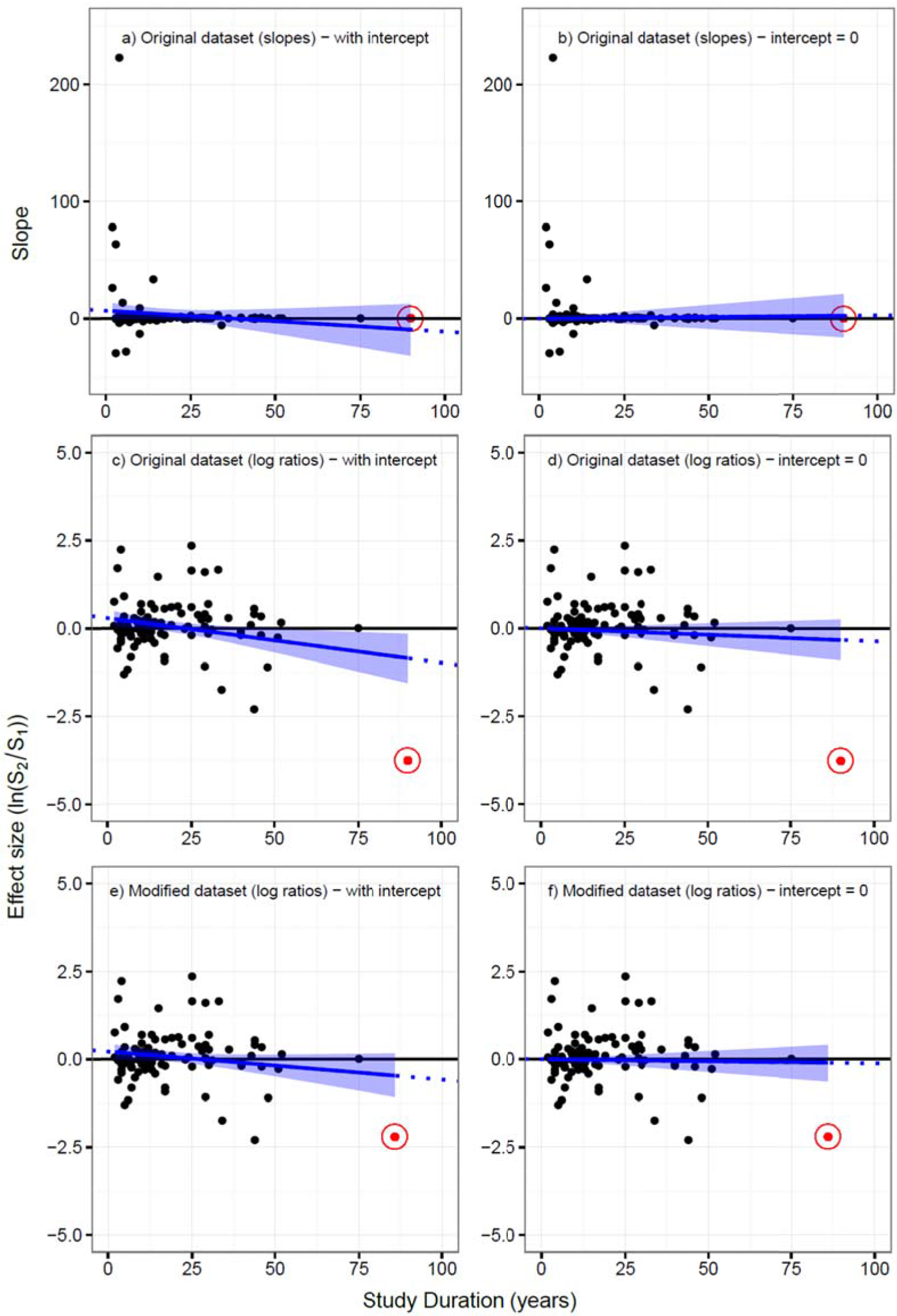
Relationships between species richness change over time (y-axis) and the duration of a study, using data from Dornelas et al. (2014). Relationships were modeled assuming either a non-zero y-intercept (a,c,e) or a zero y-intercept (b,e,f), using either slopes (a,b) or log ratios (c-f) to express temporal biodiversity change, and either including one outlier (a-d, “Original dataset”) or not (e,f, “Modified dataset”). Lines represent the estimated effect size with credible intervals. See Appendix A for statistical methods.

In sum, the evidence provided by Gonzalez et al. to support their claim that longer-duration studies tend to show biodiversity decline is exceedingly weak at best, depending on specific and unrealistic assumptions, and even then provides negligible predictive value. Whether using the realistic assumption of zero biodiversity change at duration=0, using a larger data set (always preferable), taking account of an outlier, or analyzing slopes instead of log ratios, we find no convincing evidence that estimates of biodiversity change depend on study duration. In any given time series, it is clearly possible and indeed likely that trend detection will depend on the particular period of time analyzed, but with observed trends so evenly spread above and below zero for the range of durations with lots of data (<50 years or so), there is at present no evidence to support Gonzalez et al.’s conclusion that longer-duration studies systematically show average local biodiversity declines.

### (3) The ecological literature is indeed geographically biased

Ecological studies of all kinds have been conducted far more often in Europe and North America, and nearby waters, than elsewhere. In the case of our meta-analyses, we are unable at present to estimate with confidence how local biodiversity has changed across the continent of Africa or the Indian Ocean, for example. However, while any given subset of data might deviate slightly from the overall pattern, there was no obvious signal that geographic bias led to bias against finding biodiversity decline. For example, in Vellend et al. (2013), the estimated mean log ratios of species richness change over time for South America (N = 12), Asia (N = 9), Australia (N = 5), and Africa (N = 2) were all positive. Following the selective data subsetting approach of Gonzalez et al., one could choose to conduct an analysis giving greater weight to these understudied regions: this would shift the estimated central tendency towards biodiversity increases rather than decreases. Ultimately, only new data from underrepresented regions can speak directly to what is happening in those regions, and thus prompt a potential re-assessment of conclusions. Local biodiversity change is very much dependent on specific, local circumstances, and new and interesting results from poorly known regions may well emerge in the future. Improving the spatial representation of these regions is a high priority in obtaining better estimates of local biodiversity change.

In sum, Gonzalez et al. present a sophisticated analysis to demonstrate the obvious point (noted in both original papers) that the data are geographically biased. We note that precisely the same limitation applies to most ecological synthesis and meta-analysis papers, including many by these authors (e.g., Cardinale et al. 2012, Hooper et al. 2012, Elahi et al. 2015, Haddad et al. 2015) in which there was no such vigorous effort to quantify geographic bias and its attendant consequences for limiting the scope of conclusions. In the meantime, we are working with the best data available to directly document temporal biodiversity change at the local scale. Converting natural ecosystems to croplands or parking lots causes a local loss of biodiversity (Newbold et al. 2015), but otherwise there is a great deal of variation but no clear tendency for the net temporal local biodiversity trend to be different from zero across the sites in the available data (Vellend et al. 2013, Dornelas et al. 2014, Elahi et al. 2015).

## To conclude

One point on which we agree with Gonzalez et al. concerns the need for better biodiversity monitoring in the future. Our knowledge of a great many places on earth is quite limited, and many drivers of biodiversity change are expected to push in opposite directions. For example, non-native species introductions typically increase regional-scale species richness (Sax and Gaines 2003, Winter et al. 2009), and in areas that are currently cold and humid (e.g., temperate-zone mountain tops), species richness is also expected to increase due to climate warming (Pauli et al. 2012). On the other hand, nitrogen deposition often causes plant diversity to decline (Simkin et al. 2016), and for some taxa habitat fragmentation can do the same (Haddad et al. 2015). In all of these cases, we can expect major changes in species composition-that is, species turnover-with important implications for biodiversity conservation efforts (Dornelas et al. 2014, Magurran 2016). How different forces balance out in the future can best be determined by systematic, long-term monitoring-a major priority for future research in ecology and conservation.

## Appendix A

Gonzalez et al. (in press) analysed the effect of study duration on the temporal change in biodiversity. Here we started with the same statistical models to analyse the original and updated data, and subsequently modified the models to correspond to the assumption of no diversity change for studies that last zero years (see main text).

For the Vellend et al. (2013) data, multiple data sets were recorded in many studies, which were not independent of one another. This dependency was accounted for by using a multilevel model to relate the log ratio effect sizes *y_i_* to the study duration *X_i_*, that is, *y_i_*=*α*_*j*[*i*]_ + *β*_*j*[*i*]_*X_i_* + ɛ_*i*_ (Gelman and Hill 2007). The intercepts and slopes varied between the studies according to a normal distribution with mean **μ** and standard deviation σ, so 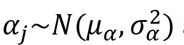 and 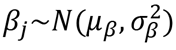. Independent within-study errors also followed a normal distribution *ε_i_~N*(0, *σ*^2^). Removing the study-level intercepts *α_j_* led to a model assuming no change in diversity for studies lasting zero years.

For the Dornelas et al. (2014) dataset, the effect sizes *y_i_* (slope or log ratio) were regressed on the duration *X_i_* with an ordinary linear model, that is, *y_i_* = *α* + *β X_i_* + ɛ_*i*_, with an intercept α, slope β and normally distributed errors *ε_i_~N(0,σ^2^)*. Again, the assumption of no diversity change for studies that last zero years was implemented by removing the intercept parameter.

The models were run in a Bayesian framework using the Stan probabilistic modelling language, called from R using the ‘*rstan*’ package (Stan Development Team 2015a). We used normal priors for the regression coefficients (α,β) and location parameters (μ_α_,μ_β_) and half-Cauchy priors for the scale parameters (σ’s) (Stan Development Team 2015b). The models were run for 5000 iterations of both warm up and sampling. Model convergence was assessed by running four chains with different starting values and by checking the trace plots and Rhat statistic. Analyses were performed in R version 3.2.4. All data and reproducible code for the analyses and graphs is available through https://github.com/lbaeten/DivChangeResponse

## LITERATURE CITED

Cardinale, B. J., J. E. Duffy, A. Gonzalez, D.U. Hooper, C. Perrings, P. Venail, A. Narwani, G.M. Mace, D. Tilman, D.A. Wardle, A.P. Kinzig, G.C. Daily, M. Loreau, J.B. Grace, A. Larigauderie, D.S. Srivastava, and S. Naeem. 2012. Biodiversity loss and its impact on humanity. Nature 486:59–67.

Colwell, R. K. and D. C. Lees. 2000. The mid-domain effect: geometric constraints on the geography of species richness. Trends in Ecology & Evolution 15:70–76.

Dornelas, M., N.J. Gotelli, B. McGill, H. Shimadzu, F. Moyes, C. Sievers, and A. E. Magurran. 2014. Assemblage time series reveal biodiversity change but not systematic loss. Science 344:296–299.

Elahi, R., M. I. O'Connor, J.E. Byrnes, J. Dunic, B. K. Eriksson, M. J. Hensel, and P. J. Kearns. 2015. Recent trends in local-scale marine biodiversity reflect community structure and human impacts. Current Biology 25:1938–1943.

Gelman, A. and J. Hill. 2007. Data Analysis Using Regression and Multilevel/Hierarchical Models. Cambridge University Press.

Gonzalez, A., B. J. Cardinale, G. R. H. Allington, J. E. Byrnes, K. A. Endsley, D. G. Brown, D. U. Hooper, F. Isbell, M. I. O’Connor, and M. Loreau. in press. Estimating local biodiversity change: a critique of papers claiming no net loss of local diversity. Ecology.

Haddad, N. M., L. A. Brudvig, J. Clobert, K. F. Davies, A. Gonzalez, R. D. Holt, T. E. Lovejoy, J. O. Sexton, M. P. Austin, and C. D. Collins. 2015. Habitat fragmentation and its lasting impact on Earth’s ecosystems. Science Advances 1:e1500052.

Hooper, D. U., E. C. Adair, B. J. Cardinale, J. E. K. Byrnes, B. A. Hungate, K. L. Matulich, A. Gonzalez, J. E. Duffy, L. Gamfeldt, and M. I. O'Connor. 2012. A global synthesis reveals biodiversity loss as a major driver of ecosystem change. Nature 486:105–108.

Magurran, A. E. 2016. How ecosystems change. Science 351:448–449.

Newbold, T., L. N. Hudson, S. L. Hill, S. Contu, I. Lysenko, R. A. Senior, L. Börger, D. J. Bennett, A. Choimes, and B. Collen. 2015. Global effects of land use on local terrestrial biodiversity. Nature 520:45–50.

Pauli, H., M. Gottfried, S. Dullinger, O. Abdaladze, M. Akhalkatsi, J. L. B. Alonso, G. Coldea, J. Dick, B. Erschbamer, R. F. Calzado, D. Ghosn, J. I. Holten, R. Kanka, G. Kazakis, J. Kollár, P. Larsson, P. Moiseev, D. Moiseev, U. Molau, J. M. Mesa, L. Nagy, G. Pelino, M. Puşcaş, G. Rossi, A. Stanisci, A. O. Syverhuset, J.-P. Theurillat, M. Tomaselli, P. Unterluggauer, L. Villar, P. Vittoz, and G. Grabherr. 2012. Recent plant diversity changes on Europe’s mountain summits. Science 336:353–355.

R Core Team (2016). R: A language and environment for statistical computing. R Foundation for Statistical Computing, Vienna, Austria.

Simkin, S. M., E. B. Allen, W. D. Bowman, C. M. Clark, J. Belnap, M. L. Brooks, B. S. Cade, S. L. Collins, L. H. Geiser, and F. S. Gilliam. 2016. Conditional vulnerability of plant diversity to atmospheric nitrogen deposition across the United States. Proceedings of the National Academy of Sciences 113:4086–4091.

Stan Development Team (2015a). Stan: A C++ Library for Probability and Sampling, Version 2.8.0.

Stan Development Team (2015b). Stan Modeling Language User’s Guide and Reference Manual, Version 2.6.1. URLhttp://mc-stan.org/.Sax, D. F. and S. D. Gaines. 2003. Species diversity: from global decreases to local increases. Trends in Ecology & Evolution 18:561–566.

Vellend, M., L. Baeten, I. H. Myers-Smith, S. C. Elmendorf, R. Beauséjour, C. D. Brown, P. De Frenne, K. Verheyen, and S. Wipf. 2013. Global meta-analysis reveals no net change in local-scale plant biodiversity over time. Proceedings of the National Academy of Sciences 110:19456–19459.

Winter, M., O. Schweiger, S. Klotz, W. Nentwig, P. Andriopoulos, M. Arianoutsou, C. Basnou, P. Delipetrou, V. Didžiulis, and M. Hejda. 2009. Plant extinctions and introductions lead to phylogenetic and taxonomic homogenization of the European flora. Proceedings of the National Academy of Sciences 106:21721–21725.

